# A heterochromatic knob reducing the flowering time in maize

**DOI:** 10.1101/2021.03.31.437909

**Authors:** Renata Flávia Carvalho, Margarida Lopes Rodrigues Aguiar-Perecin, Wellington Ronildo Clarindo, Roberto Fristche-Neto, Mateus Mondin

## Abstract

Maize flowering time is an important agronomic trait, which has been associated with variations in the genome size and heterochromatic knobs content. We integrated three steps to show this association. Firstly, we selected inbred lines varying for heterochromatic knob composition at specific sites in the homozygous state. Then, we produced homozygous and heterozygous hybrids for knobs. Second, we measured the genome size and flowering time for all materials. Knob composition did not affect the genome size and flowering time. Finally, we developed an association study and identified a knob marker on chromosome 9 showing the strongest association with flowering time. Indeed, modelling allele substitution and dominance effects could offer only one heterochromatic knob locus that could affect flowering time, making it earlier rather than the knob composition.

## INTRODUCTION

The relationship between heterochromatic knobs, genome size, and flowering time (FT) in maize is a long debate. Knobs have intrigued geneticists for more than 100 years, and since their discovery, their functions remain under investigation. These maize genome regions are constitutive heterochromatin with late replication during the cell cycle (Pryor et al., 1980), extensively composed of two highly repetitive satellite DNA families, 180-kb, and TR-1 (Peacock et al., 1981; Ananiev et al., 1998). An eminent observation was their wide variability among landraces, inbred lines, and hybrids, following a biogeographic distribution (McClintock et al., 1981; Rayburn et al., 1985), suggesting a possible role affecting the expression of some phenotypic features as flowering time (Jian et al., 2017).

Moreover, other exciting aspects of the heterochromatic knobs in the maize genome have been elucidated by several studies. Knobs have been shown to affect local recombination (Ghaffari et al., 2013; Stack et al., 2017) and genes adjacent to these regions reduce their expression level (Haberer, 2020). The meiotic drive mechanism influences the number of knobs in different species of *Zea* (Rhoades and Dempsey, 1966; Buckler et al., 1999; Higgins et al., 2018). Such a system would favor knob transmissions preferentially during female meiosis by a knob neocentromere activity, which might have contributed to maize genome remodeling throughout its evolutionary history. Response to abiotic stress was observed in maize plants from the transcriptional activation of the knob repetitive sequences, being this activation selective, temporary, and accompanied by epigenetic changes (Hu et al., 2012). Lastly, associations between heterochromatic knobs and agronomic traits have also been reported (Blumenschein, 1964).

There is a growing interest in the role of variation of genomic content in creating phenotypic modifications within a species. These changes are due to copy number variation (CNV), which has been used to describe duplications, deletions, and insertions in individuals of a species, and presence/absence variation (PAV) that describes the presence or not of sequences on the genome of different individuals of the same species. Together, they form the pan-genome of species (Springer et al., 2009). Maize is also a model species for studies of pan-genome. Differences in chromosomal structure between maize landraces were firstly identified through cytogenetic studies. Barbara McClintock and colleagues analyzed the content and size of heterochromatic knobs to characterize this variation in the genome (McClintock et al., 1981). Currently, modern cytogenomic techniques have sampled the wide variation of the copy numbers resulting from repetitive sequences, which make up most of the maize genome (Kato et al., 2004; Albert et al., 2010; Mondin et al., 2014; Bilinski et al., 2018)

Moreover, quantification of DNA content through flow cytometry has documented significant variability in the maize genome size (GS) between landraces and inbred lines (Laurie and Bennett, 1985; Realini et al., 2015). New combinations of alleles arise from the variation of genomic content within species, contributing to phenotypic variation (Brohammer et al., 2018). Therefore, this can have an influence on several important characteristics, including flowering time (Bilinski et al., 2018). Flowering time is a quantitative trait of extreme relevance for cultivated plants since it controls plant adaptation to the environment. Breeding programs through this feature outline strategies to make earlier or later varieties, allowing their expansion to other regions and consequently increasing yield (Jung and Müller, 2009).

Recent studies on maize genome present the triangle formed between genome size, knob content, and flowering time in natural populations. These surveys have considered just the knob numbers or estimated the knob abundances by low coverage sequencing (Realini et al., 2015; Jian et al., 2017; Bilinski et al., 2018). However, few studies have evaluated the knob effect depending on its homozygous or heterozygous condition (Chughtai and Steffensen, 1987). These authors reported that heterozygotes for knobs had earlier flowering time in comparison with homozygotes. Different researchers have attributed variation in maize genome size to heterochromatin content differences, especially in knobs (Chughtai and Steffensen, 1987; Chughtai et al., 1997; Jian et al., 2017) However, knob content not corresponding with genome size was also reported (Realini et al, 2015). Moreover, it is proposed that there is a relationship between decreased genome sizes in high altitudes with reduced flowering time. Besides, the same association is observed for knob abundance occurring along altitudinal clines (Poggio et al., 1998; Bilinski et al., 2018). However, no experimental design has isolated the effects of homozygous or heterozygous knobs and test whether they are correlated with flowering time. In this case, the relationship between knobs and flowering time remains unclear.

Here, we derived a panel of sister inbred lines and their hybrids to verify the association between genome size, knob constitution, and flowering time for male (MF) and female (FF) inflorescences. Firstly, in chronological order, inbred lines with variable presence or absence of the knobs K3L, K5L, K7S, and K9S were derived until S9 generation from a segregating single seed of a flint variety (Decico, 1991). Then, we created hybrids between inbred lines, and finally, we started to estimate the genome size and flowering time.

These near-isogenic inbred lines and their hybrids, varying for knob positions at K3L,K5L, K7S, and K9S were used to assay flowering time under controlled environmental conditions and their genome size was measured by flow cytometry. All data were used to analyze the genome association study. We expected that using sister inbred lines and their hybrids possessing a common genetic background, the influence of knobs on genome size and flowering time could be clearly detected (Carvalho et al., 2021).

## MATERIALS AND METHODS

### PLANT MATERIAL

#### Origin of the maize inbred lines and knob composition

For this research, the development process of inbred lines was idealized in the Department of Genetics at “Luiz de Queiroz” College of Agriculture – ESALQ, University of São Paulo – USP. The initial biological material was a commercial variety Jac-Duro (JD – composed of Cateto varieties, Flint type endosperm), donated by Agroceres Seeds, Brazil. Initially, the variety segregated for the knob positions K3L, K5L, K7S, K9S (where “K” refers to a knob, number corresponds to chromosome, L or S is for long and short arm, respectively), and it was homozygous for knob positions at K6L2, K6L3, K7L, and K8L1, KL2. Thus, over the self-fertilization cycles, the inbred lines were analyzed by C-banding until they become homozygous for specific knob positions. (Table S1). The knobs K6L2 and K6L3, and K8L1 and K8L2 visualized at pachytene are seen as a unique band in somatic chromosomes using C-banding or FISH (Mondin et al., 2014). Analyses of the C-band frequency in somatic chromosomes were performed on S2 (240-14-4) and S3 (240-14-1 and 240-14-2) until S9 progenies (Decico, 1991). The S3 240-14-1 progeny segregated for two knob positions: K3L and K9S, and progeny 240-14-2 for one locus: K9S. The S2 240-14-4 progeny segregated for the four loci: K3L, K5L, K7S, and K9S. All the inbred lines maintained the homozygous knobs for the positions K6L2, K6L3, K7L, and K8L1 and K8L2. S4 and S5 inbred lines were obtained from these progenies, derived from two self-fertilization cycles that became denominated in S5 = 14-1-3 and 14-2-1 and S4 = 14-4-1 and 14-4-4. This designation was abbreviated for JD 1-3, JD 2-1, JD 4-1, and JD 4-4 to characterize the four families of lines. The inbred lines used in this research were derived until S9 progenies (Table S1) (Decico, 1991), and the hybrids were obtained from crosses between JD 1-3 and JD 4-4 inbred lines (Table S2). Therefore, we have a broad panel of heterochromatic knob combinations, which can be homozygous for the presence or absence or even heterozygous in the hybrids. Figure 1 illustrates these conditions and Figure 2 shows a sample of the inbred lines and hybrids. Mondin et al. (2014) described the karyotype of these lines through the analysis of pachytene and somatic chromosomes using C-banding and FISH procedures.

**Figure 1.**
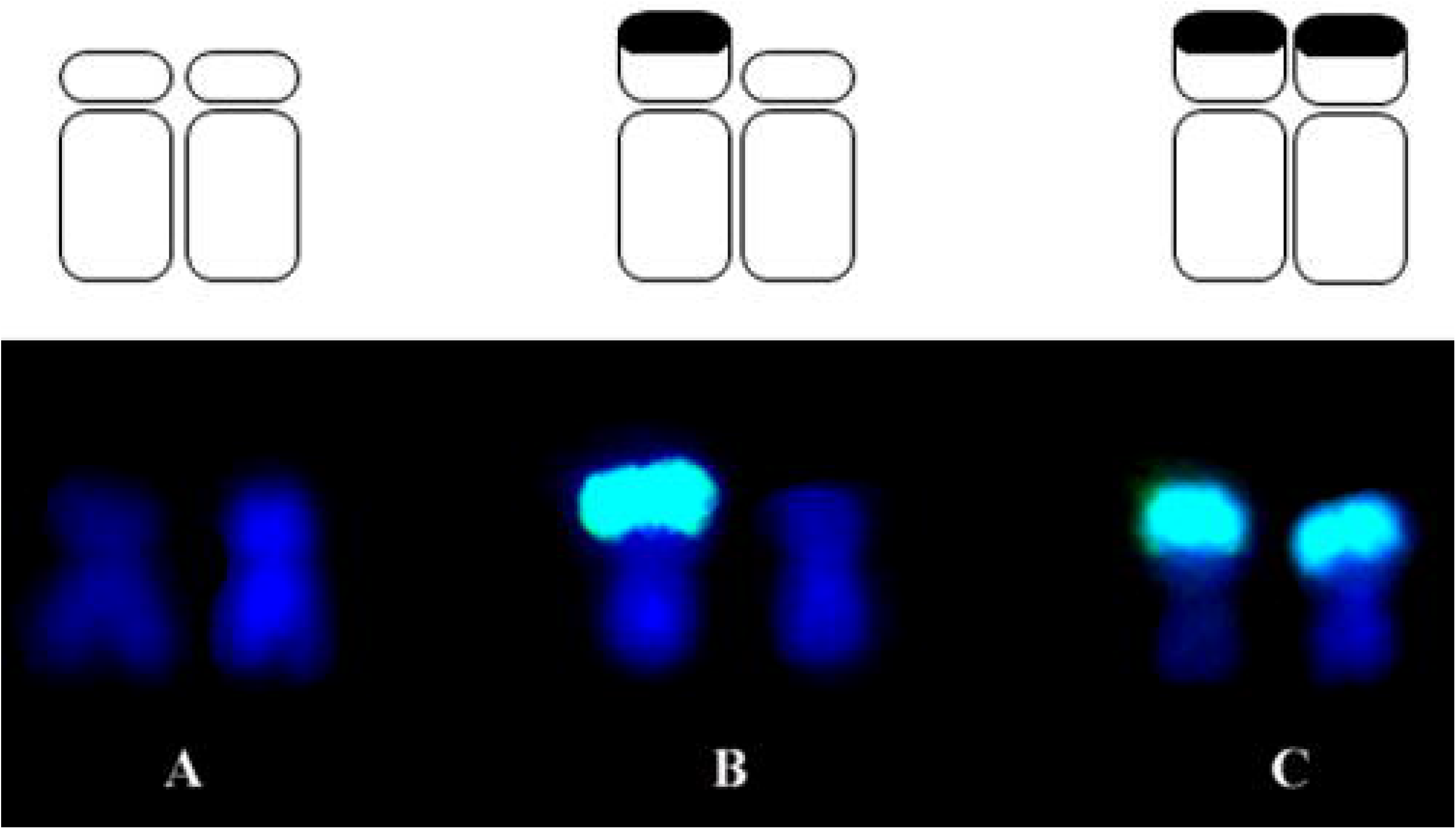
Representation of the knob conditions for chromosome 9 (K9S). **(A)** Absence of knob; **(B)** heterozygous knob and **(C)** homozygous knob.

**Figure 2.**
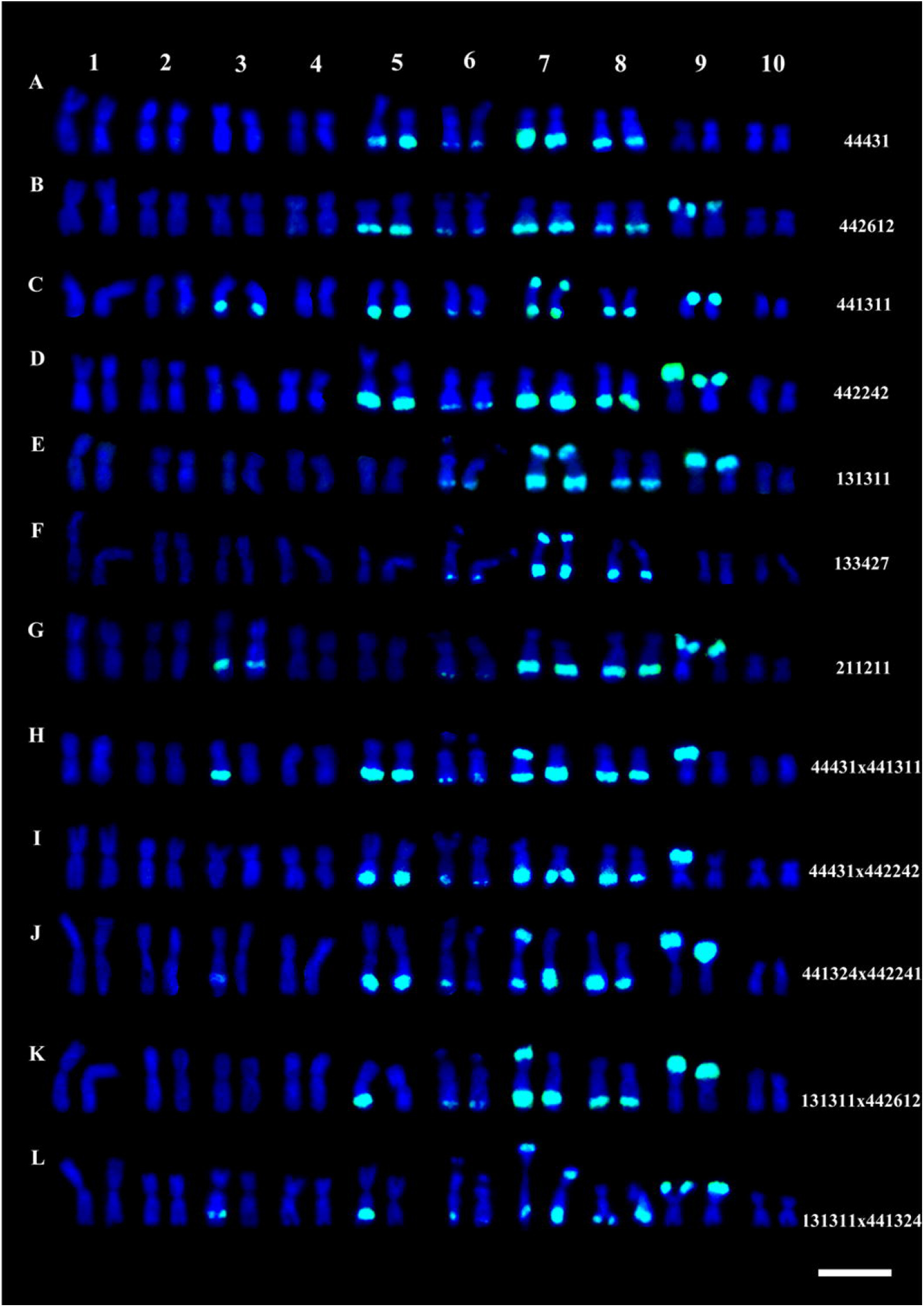
A sampling of genotypes used in the experiments on flowering time and genome size. (A) – (G) Somatic karyotypes of JD lines; and (H) – (L) hybrids, labeled by FISH using a probe for the knob 180-bp repeat (green).

## METHODS

### Fluorescence *in situ* hybridization - FISH

The 180 bp repetitive DNA sequence probe present in the knobs (Peacock et al., 1981) was used to map the inbred lines and hybrids. The steps of pre-treatment and *in situ* hybridization were based on Mondin et al. (2014). Each chromosome preparation was conducted using 20 μl of a probe mixture containing the 180 bp probe. The probe mixture was denatured by heating at 96ºC for 10 minutes, cooled on ice, and then dropped onto the slide preparations and covered with a coverslip. Preparations were denatured in a thermocycler at 93ºC for 10 minutes. The hybridization was performed at 37°C for 16h. Following the hybridization, slides were washed twice in 2x SSC at 37ºC and 42ºC for 5 minutes, twice in 20% formamide in 0.5 × SSC at 42ºC (74% stringency) for 10 minutes and once in 0.5x SSC for 5 minutes at the same temperature. The probe of the 180 bp knob repeat was directly labeled with 6-Carboxyfluorescein (FAM). The slides were counterstained with 0.2 μg/ml 4,6-diamidino-2-phenylindole (DAPI) and mounted in 5 μl of Vectashield H-1000.

### Genome size measurements

The genome size of the inbred lines and hybrids was estimated following Praça-Fontes et al. (2011). For these analyses, *Z. mays* ‘CE-777’ was used as an internal standard. Young leaves of the sample and standard were chopped into a Petri dish containing a solution of 0.5 ml of OTTO I nuclear extraction buffer (Otto, 1990). To this solution it was added 2.0 mM of dithiothreitol and 50 µg/ml of RNase. Later, it was added the same volume of buffer solution. This homogenate was filtered, centrifuged for 10 minutes and resuspended in OTTO I buffer. The samples were stained in 1.5 ml of OTTO-I:OTTO-II (1: 2) staining buffer (Otto, 1990), supplemented with 50 mM dithiothreitol, 50 µl RNAse, and 75 µM propidium iodide, for 20 minutes, to define the size of the nuclear genome (Dolezel et al.et al., 2007). Five replicates were used for each sample.

Nuclear suspensions were analyzed in a Partec PAS® flow cytometer (Partec® Gmbh, Munster, Germany), equipped with a laser source (488 nm) and a UV lamp (388 nm). The histograms were used to measure the nuclear genome by comparing fluorescence peaks corresponding to the G_0_/G_1_ stages of the standard (CE 777) and samples (inbred lines and hybrids). The genome size measurements were performed at the Laboratório de Pesquisa em Citogenética e Citometria, at the Federal University of Viçosa.

### Experimental design to analyze flowering time

Two experiments were conducted in a greenhouse under 28ºC/25ºC day/light and 12h light/12h dark in two subsequent years (2018/2019), for the analysis of flowering time. The first assay counted with 8 inbred lines (parents) and 35 hybrids. The experiment was carried out in a completely randomized design, with 3 replicates for parents (24 plants) and 5 replicates for hybrids (175 plants). The second assay was performed with 20 inbred lines with 5 replicates each one, totalizing 100 plants. Both experiments were conducted from February to June, and the maize plants were planted in 20L pots with 50 cm spacing between them. Flowering time was calculated as the number of days from planting until the first day of flowering. Inbred lines and hybrids with different knob constitutions were evaluated individually for male flowering (MF) and female flowering (FF).

### Association study

A Mixed Linear Model (MLM) was run by the FarmCPU R package (Liu, 2016) to determine the knob-trait associations. The MLM equation used in the analysis was as follows:

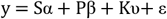

where: *y* is the BLUP of the genotypes for AN or ALS; α is the vector of fixed effects of the knobs; β is the vector of fixed effect of the population structure (first principal components used, depending on the trait); υ is the random effect of the relative kinship, where 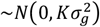.; *ε* is the error term, where 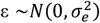. S, P, and K are incidence matrices that relate the independent vector effects from each matrix with the dependent *y* vector.

The additive and heterozygous (dis)advantage models were applied in adapted GWAS analyses using specifics encodings for the knob matrix (in this case, replacing SNPs). Heterochromatic knobs present a Mendelian inheritance pattern; therefore, individuals could be homozygous for knobs, homozygous for absence, or heterozygous for knobs (Aguiar-Perecin and Decico, 1988). This feature can be used analogously to the SNP marker, which is co-dominant bi-allelic, precisely as the knobs behavior. Concerning the additive knob effect with two alleles (A1 and A2), the knob matrix was coded by 0 (A1A1), 1(A1A2), and 2 (A2 A2),considering the A2 as the minor allele or knob absence. In this context, the additive GWAS model assumes a linear change in the phenotype regarding the minor allele number of copies. On the other hand, in the heterozygous (dis)advantage GWAS model,the homozygous genotypes (A1A1 or A2A2) were assumed to have the same effect. In contrast, the heterozygous genotypes have a different one, implying an increase or decrease in the trait effect. Therefore, the knob matrix was coded by 0 (A1A1), 1 (A1A2), and 0 (A2A2) (Tseplov et al., 2015).

To determine the *p*-value threshold, we used a resampling method. Therefore, first, the phenotypic values are shuffled, breaking their association with markers, and then the random association between all markers to the phenotype is estimated, and the corresponding best marker score (minimum *p*-value obtained among all markers) is recorded. This procedure was repeated 50 times for each trait, and the 95% quantile from the 50 best scores was defined as the threshold to declare a significant association.

## RESULTS

### KNOB COMPOSITION AND MAIZE GENOME SIZE

The genome size of the inbred lines and hybrids was measured to test whether knob composition contributes to the DNA content variation. Table S2 shows the values of genome size and knob composition of the materials. Each inbred line and hybrid can have different knob configurations at the K3L, K5L, K7S, and K9S positions on the chromosomes. These combinations can be homozygous for presence or absence of knobs in the lines or heterozygous in the hybrids (Figures 1 and 2). Note that there are hybrids homozygous and heterozygous for knobs.

The genome size was considered a quantitative trait, and its heritability was estimated at 26%. The hybrids presented a higher mean genome size than inbred lines, showing 5477 Mbp (2C = 5.6 pg) and 5281 Mbp (2C = 5.4 pg), respectively (Fig. 3). However, no significant differences were found. Even when homozygotes were compared with heterozygotes, no differences were observed (Fig. S1).

**Figure 3.**
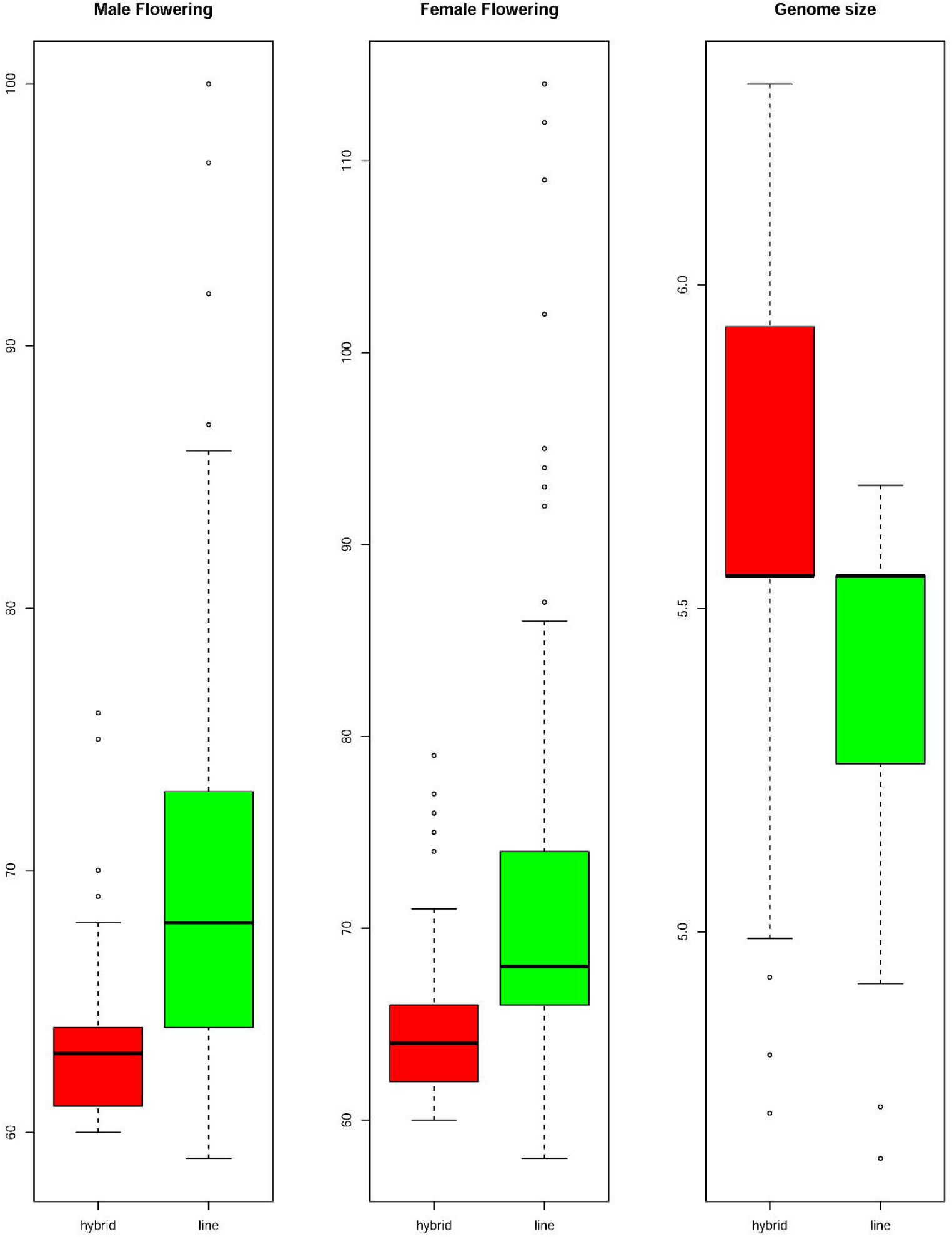
Flowering time and Genome Size of the hybrid and inbred lines. Boxplots compare mean values between male and female flowering time and genome size for inbred lines and hybrids. The flowering time is showed in days to flowering and the genome size in picograms. The boxes indicate the first quartile (lower line), the second quartile or mean (central line), and the third quartile (upper line). Additionally, the whiskers represent the standard deviation with the dots as the outliers.

The results showed that knob dosages are not enough to explain the genome size increase or decrease in the inbred lines and hybrids. However, genome size varied broadly, despite the same knob composition. For instance, the hybrids presenting the highest (441311/2 × 441324/1, 2C = 6.31 pg) and the smallest genome size (442213/1 × 441311/2, 2C = 4.72 pg) share the same knob constitution (Table S2). For inbred lines, the largest genome size (131311/1-04, 2C = 5.69 pg) has a lower number of knobs than the shortest genome (442213/1, 2C = 4.65 pg), which is homozygous for knob presence in the four positions described (Table S2).

An adapted genome-wide association study (GWAS), with the heterochromatic knob full panel to identify associations with genome size and flowering time, was performed, where the knobs were used as genetic markers. Our null hypothesis was that there were significant differences in genome size and days for male and female flowering due to the knob conditions in the K3L, K5L, K7S, and K9S positions. To test our hypothesis, we used the inbred lines and hybrids panel, to which all knobs were mapped. The panel has a matrix-*like* structure with different knob combinations for presence (++ or 1) or absence (00 or -1) when homozygous and as heterozygous when just one of the homologous has a knob (+0 or 0). Firstly, only the allele substitution effect model was performed, and no significant association was found between knobs and genome size (Fig. S2). Moreover, this information was supported by a null correlation between genome size and knob dosage classes (Table 1).

**Table 1.** Pearson’s correlation between traits and knobs.

|  | Male flowering | Genome size | Knob dosage classes |
| --- | --- | --- | --- |
| Female flowering | 0.95 | -0.14 | -0.20 |
| Male flowering |  | -0.17 | -0.24 |
| Genome size |  |  | -0.08 |

### GENOME SIZE AND KNOB CONSTITUTION DID NOT AFFECT MAIZE FLOWERING TIME

Days to MF and FF were evaluated individually (Table S2) and heritability was estimated to be 51% for MF and 41% for FF. The mean values for days to MF and FF for hybrids were 63 and 64 days, respectively, while the FT for inbred lines was 70 days for MF and 72 days for FF (Figure 3). Heterochromatic knob configurations as heterozygotes exhibited shorter flowering times than those in homozygous states (Figure S1). The flowering time data were also plotted showing its amplitude inside the inbred line families and hybrids (Figure 4).I t was observed synchronicity for both traits (MF and FF) within each group analyzed, and hybrids had shorter flowering time than inbred line families.

**Figure 4.**
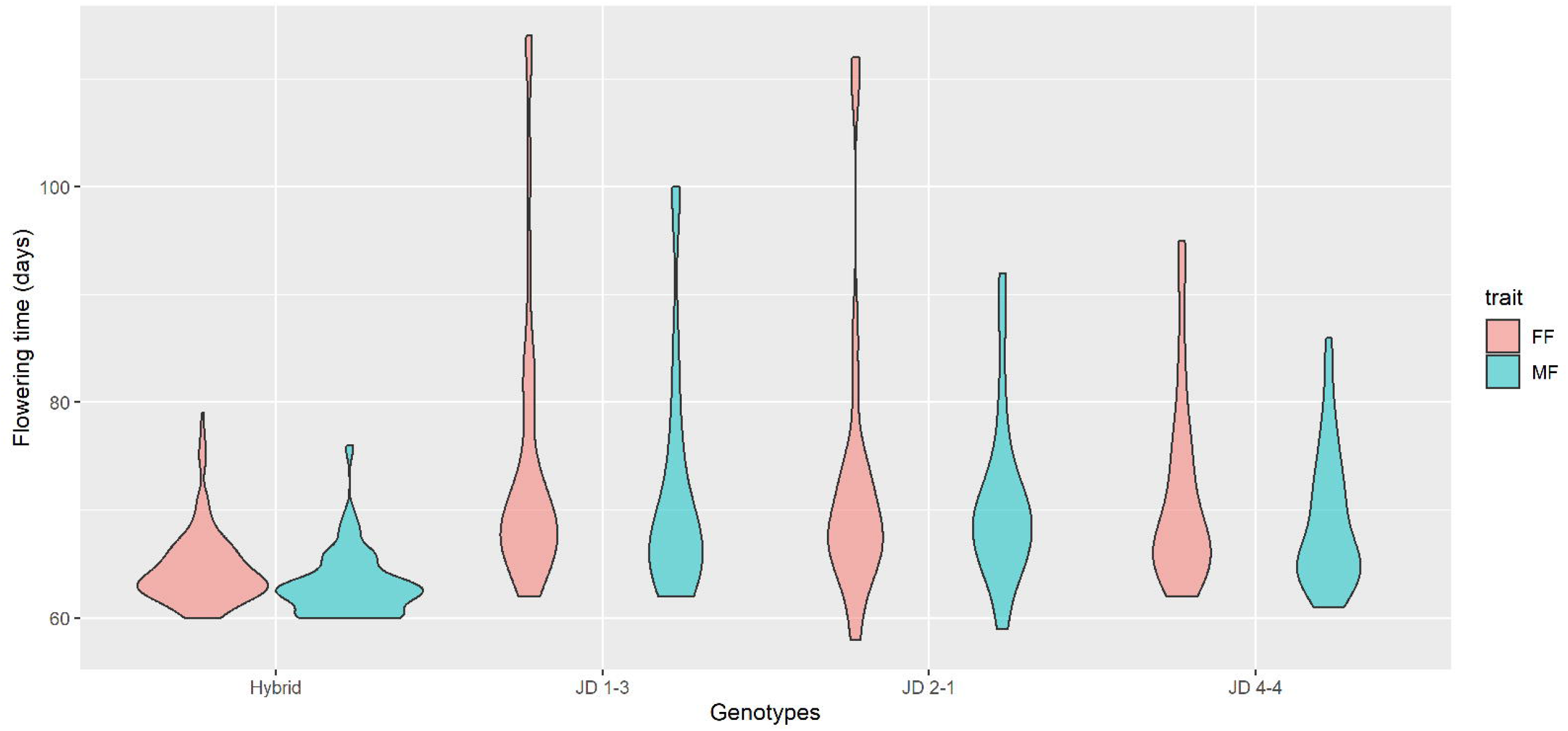
Flowering time among inbred line families and hybrids. The violin plot shows the FT distribution in days for all analyzed materials. The violin shape represents the estimated value of the density of the trait within each group. Hybrid, n = 175; JD 1-3, n =36; JD 2-1, n = 25; JD 4-4, n = 63.

Further, the flowering time amplitude of hybrids was much smaller than the other groups, ranging mainly between 60 and 70 days, in agreement with the observed mean values. A broader variation in flowering time was observed for inbred line families (58 - 114 days), with distribution densities mainly varying between 60 and 80 days. The JD 4-4 family had a smaller flowering time distribution for FF and FM among families.

Negative values for Pearson’s correlation were found comparing female and male flowering time with knob dosage classes (Table 1). The knob combinations were used to create dosage classes and test their effects on genome size and flowering time. The knob dosage classes were not correlated with the flowering time, i.e., an increase in knob numbers did not correlate with late flowering time. It was possible to observe that the mean values of MF and FF were very similar for each knob dosage class (Figure 5). It was also interesting to note that the inbred lines with only two knobs (dosage class = 2) flowered later than those with other dosage classes (dosage class = 3-8). Figure 5 also shows that the knob dosage classes did not alter the genome size.

**Figure 5.**
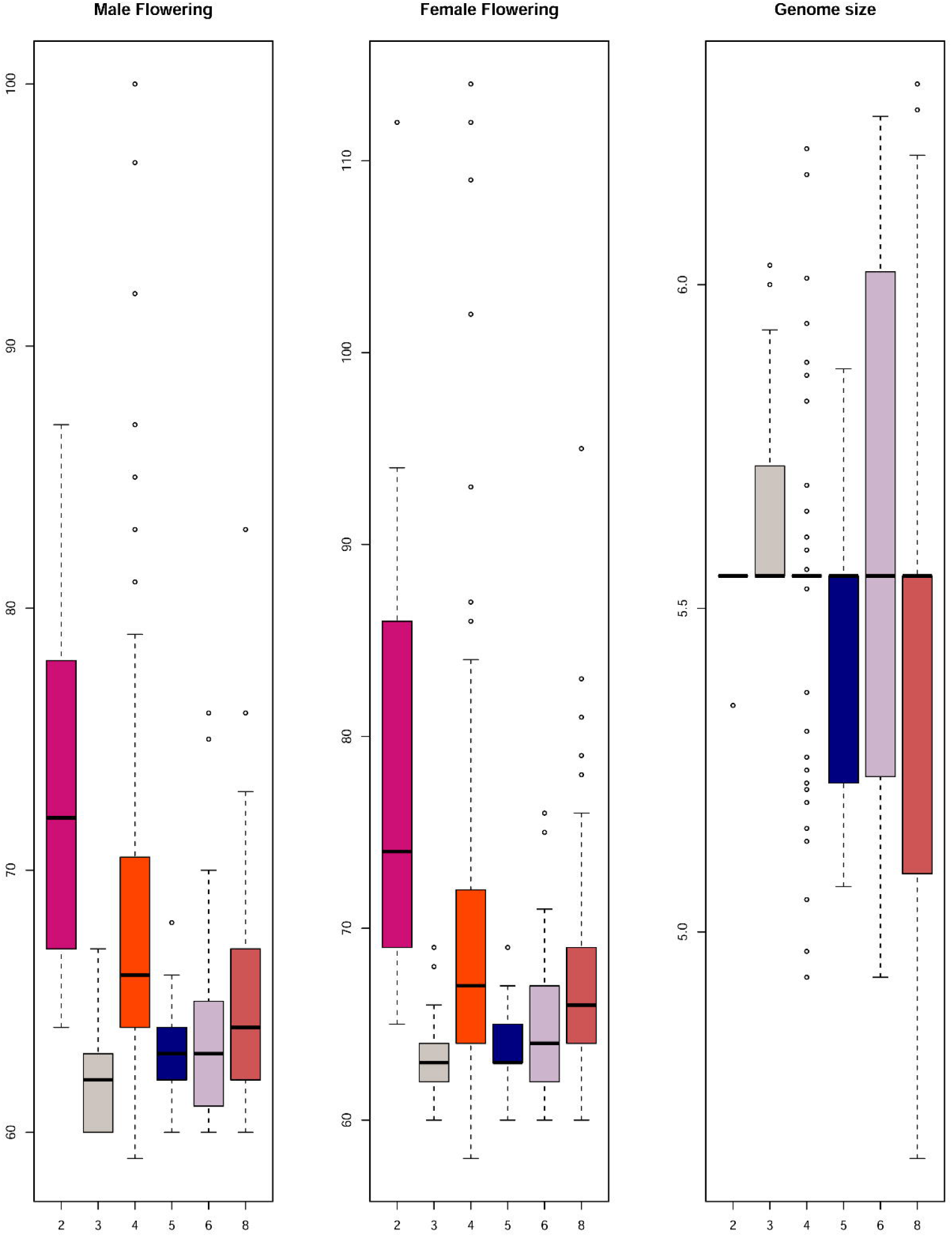
Comparison between dosage classes of knobs and male and female flowering time, and genome size. The flowering time is expressed in days to flowering and genome size in picograms (y axis). The classes mean all possible combinations of presence and absence, homozygous or heterozygous, of the knobs considering only chromosomes 3, 5, 7 and 9 (x axis).

Furthermore, two GWAS models to illustrate the interactions between flowering time and knob conditions were used. The first model took into account only the allele substitution effect of the markers (Figures 6A, 6B, and S3), and the second considered the dominance effect (Figures 6C, 6D, and S4). This analysis showed only one significant marker-trait association for both allele substitution and dominance effect models regarding flowering time. This significant association was observed just for the knob on the short arm of chromosome 9 (K9S) (Figure 6). Regardless of the GWAS model, the knob marker on chromosome 9 displayed the same performance concerning the flowering time, showing the *p*-value highly significant (Table 2), while for the knobs K3L, K5L and K7S the p-value was not significant for flowering time and genome size. (Table S3).

**Table 2.**
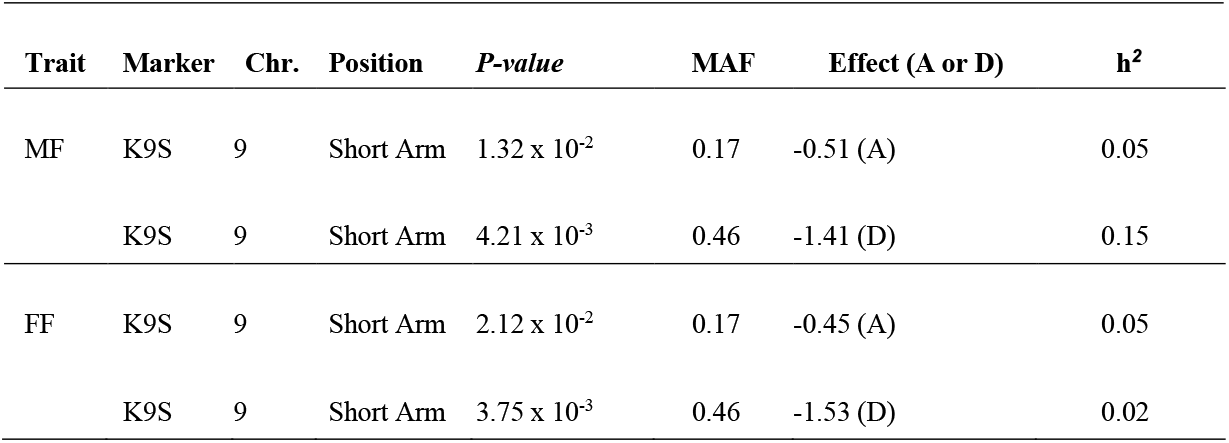
Significant marker-trait association for flowering time. Male flowering (MF), female flowering (FF), knob in the short arm of chromosome 9 (K9S), *p-*value Bonferroni test, minor allele frequency (MAF), allele substitution model (A), dominance model (D), and heritability (h^2^).

| Trait | Marker | Chr. | Position | *P*-value | MAF | Effect (A or D) | $h^2$ |
| --- | --- | --- | --- | --- | --- | --- | --- |
| MF | K9S | 9 | Short Arm | $1.32 \times 10^{-2}$ | 0.17 | -0.51 (A) | 0.05 |
| | K9S | 9 | Short Arm | $4.21 \times 10^{-3}$ | 0.46 | -1.41 (D) | 0.15 |
| FF | K9S | 9 | Short Arm | $2.12 \times 10^{-2}$ | 0.17 | -0.45 (A) | 0.05 |
| | K9S | 9 | Short Arm | $3.75 \times 10^{-3}$ | 0.46 | -1.53 (D) | 0.02 |

**Figure 6.**
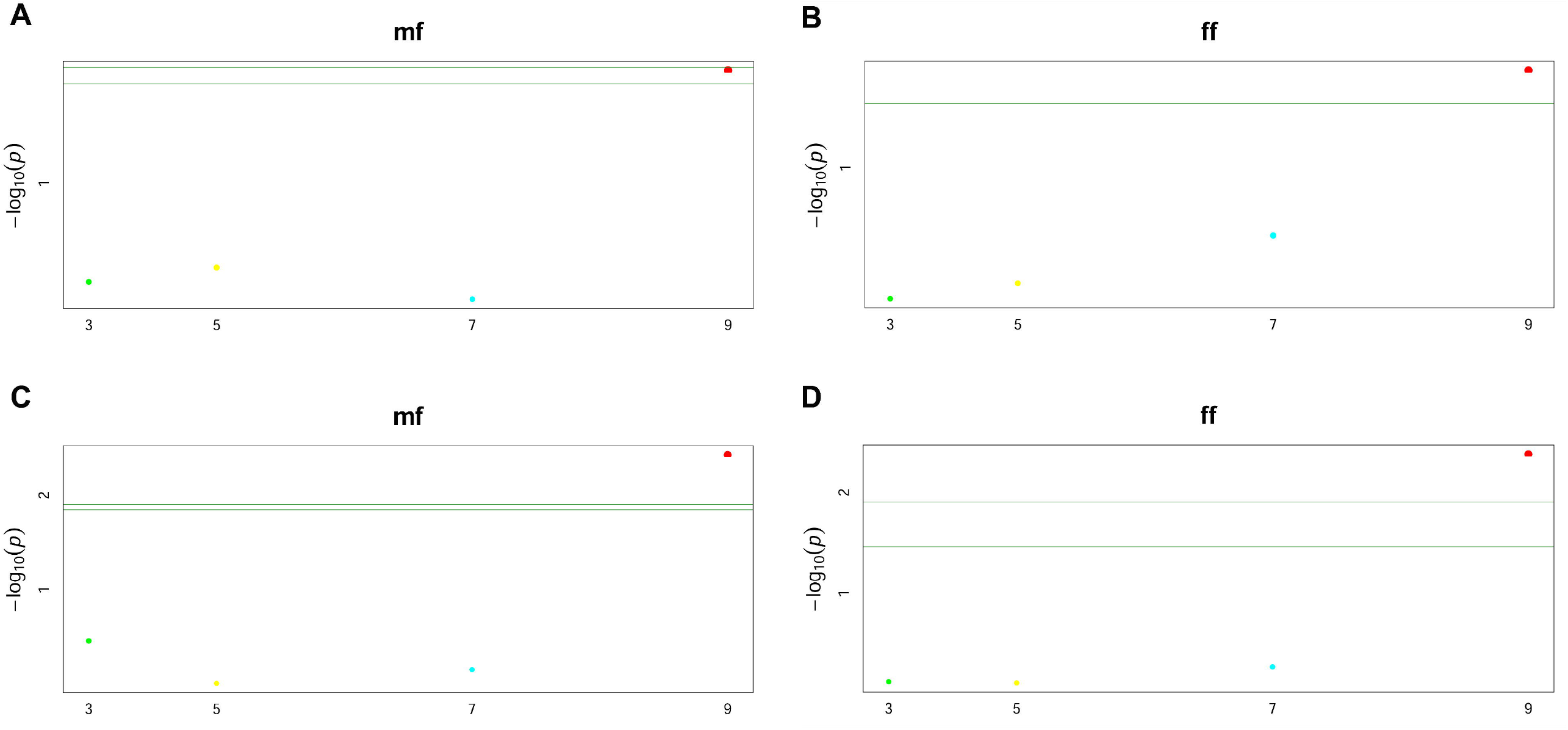
Manhattan plots of GWAS for male (mf) and female (ff) flowering time using the knob positions K3L, K5L, K7S, and K9S. (A) and (B) are plots showing the allele substitution effect model, (C) and (D) the dominance effect model, both with p-values < 0.05. The green lines are the significance threshold, a Bonferroni-corrected significance threshold used to identify significant associations.

Regarding the allele substitution effect model, the presence of the K9S decreased the MF and FF of the inbred lines and hybrids by -0.45 and -0.51 days, respectively. While on the dominance effect model, this effect was even more significant, decreasing flowering time in one and a half days (MF = -1.40 and FF = - 1.53) (Fig. 7). Only the knob on chromosome 9 significantly reduced the flowering time in the inbred lines and hybrids. Decreasing in flowering time was affected by the presence of the knob in the homozygous or heterozygous configuration. The heritability of the marker for both models varied from 0.02 to 0.15 (Table 2). Our analysis also showed slight effects of the knobs K3L, K5L and K7S on flowering time, albeit these were not significant (Table S3). These results showed that the knob composition was not essential to model the genotype effects on the trait. The flowering time violin plot (Figure 4) also supported the GWAS analyses (Fig. 6), since all hybrids are homozygous or heterozygous for the K9S, corroborating the specific contribution of this knob to decrease maize flowering time.

**Figure 7.**
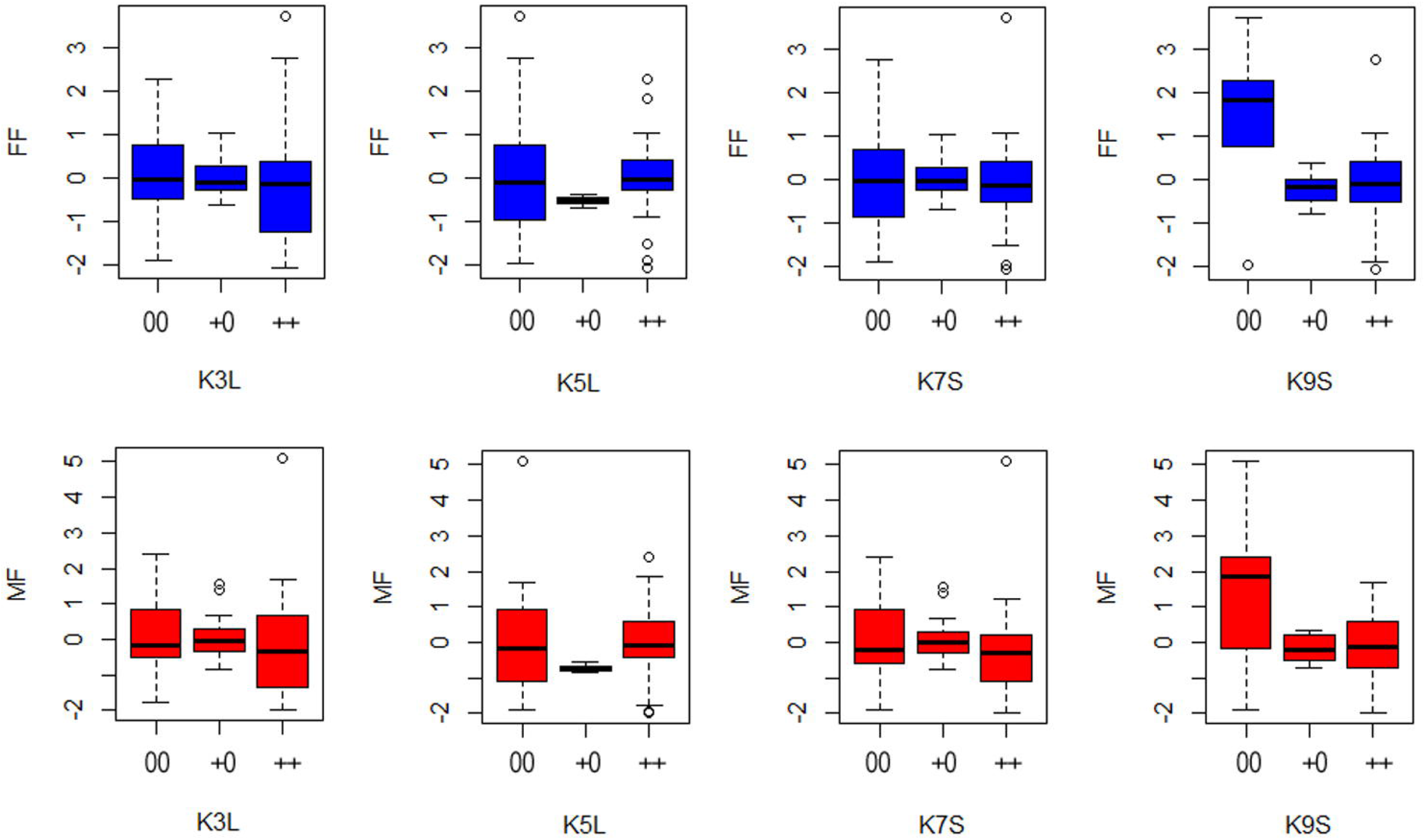
Boxplots show male (MF) and female (FF) flowering time for each knob condition in the K3L, K5L, K7S, and K9S positions, considering both allele substitution and dominance effects. The x-axis shows the classification for each knob condition 00 = knob absence; +0 = heterozygous knob presence and ++ = homozygous knob presence. The y-axis shows the genetic effects of flowering time for female and males plants.

## DISCUSSION

### KNOB COMPOSITION DID N0T CORRELATE WITH MAIZE GENOME SIZE

In our study no pattern was observed, which indicates that there is probably a linear relationship between the number of knobs present in the lines and hybrids with the increase or decrease in the DNA content. Interestingly, the hybrids presenting the highest and the smallest genome size share the same knob constitution. For inbred lines, the largest genome size has a lower number of knobs than the shortest genome size, which is homozygous for knob presence in the four positions described.

There is a wide variability of genome size in natural maize populations. The maize genome size associated with knob numbers and with flowering time is reported in some studies. Most of these studies have shown positive correlations between genome size and knob content (Rayburn et al., 1985; Tito et al., 1991; Jian et al., 2017; Forastié et al. 2018). However, in some studies, a positive correlation between genome size and knob composition was not found (Laurie and Bennett, 1985; Realini et al., 2015). A recent study provided evidence that natural selection plays a substantial role in reducing the maize genome size at high altitudes (Bilinski et al., 2018). The authors also showed that the abundance of transposable elements (TEs) and heterochromatic knobs are significantly correlated with altitude, and the knobs act as significant effect loci in the genome size. Unlike what they found, our data did not show this same relationship.

Our results corroborate the surveys showing no significant correlation between maize population genome size and heterochromatin percentage. One hypothesis is that as our materials differ in the composition of only four knob positions, measurements by flow cytometry would not detect slight differences in genome size due to knobs. Furthermore, we might infer that differences in the genome size among our inbred lines and hybrids may result from transposable element (TE) variability, once the loss of TE during selfing has been demonstrated (Roessler et al., 2019). Knobs are additional material in the chromosomes and alter the length of somatic chromosome arms (Aguiar-Perecin and Vosa, 1985), therefore they should increase the genome size, but this effect could be masked by TE variability.

### GENOME SIZE AND KNOB COMPOSITION DID NOT AFFECT MAIZE FLOWERING TIME

The genetic architecture of flowering time in maize has been widely studied, given that this trait reflects the plant adaptation to the environment. Since maize is distributed throughout America and is adapted to a wide range of environments, understanding how flowering time is regulated is of paramount importance and generates valuable information for breeding programs (Buckler et al., 2009). Besides this study, in the last two decades, the maize flowering time was dissected using different approaches such as linkage and association mapping (Chardon et al., 2004; Ducrocq et al., 2009; Li et al., 2016; Alberto et al., 2017), population genetics (Wang et al. 2017; Guo et al., 2018), archaeological DNA studies (Yang et al., 2019), genome-wide association studies (Hung et al., 2012; Yang et al., 2013; Jian et al., 2017; Liang et al., 2019), and gene analysis (Lazakis et al., 2011; Alter et al., 2016; Minow et al., 2018; Stephenson et al., 2019)). In inbred maize lines, the genetic architecture of flowering time has been attributed to the cumulative effect of numerous quantitative trait loci (QTL), each with a small impact on this trait (Buckler et al., 2009).

Besides these reports on the genetic control of maize flowering, a positive relationship between genome size and flowering in maize have been reported (Rayburn et al., 1985; 1994; Tito et al., 1991). In addition, significant positive correlations between genome size, knob abundance, and flowering time were found in maize inbred lines (Jian et al., 2017). However, in this study, these correlations were lost when a kinship matrix was introduced in the analyses. The authors performed an association study where genome size was also correlated with flowering time. Three genomic regions associated with genome size were found, and mapped close to the knob region on chromosome 8 (Jian et al., 2017). In contrast, another study analyzing maize landraces in northern Argentina found no correlation between genome size and flowering time (Realini et al., 2015). Nevertheless, in this study a positive relationship between the vegetative cycle and heterochromatin percentage was observed. Another report argued that repetitive sequences would have indirect effects on flowering time due to their impact on genome size and might depend on the environment (Bilinski, 2018).

However, this effect was not found in our study since there was no relationship between genome size, knob composition and flowering time. It is interesting to note that the heterozygotes had an early flowering time in comparison with lines. This result is in agreement with the findings by Chughtai and Steffensen (1987) that also showed a positive correlation between knob content and flowering time.

In fact, the relationship between genome size and phenotype in plants has been reported (Greilhuber and Leitch, 2013). Especially, the genome size has been correlated with the duration of the cell cycle and this feature could affect the vegetative cycle. We could argue that as knobs have late replication in the mitotic cycle, they would replicate simultaneously, and then, differences in their number would not affect the duration of the cell cycle. However, if we compare populations differing in the size of knobs, we could suppose that the larger knobs would increase the duration of the mitotic size. As in our study we compared lines differing in few knobs approximately with the same size, differences in flowering time were not observed.

Another kind of studies had already indicated that gene groups responsible for the plant morphological and physiological traits would be correlated with the presence of knobs (Blumenschein, 1964; Rhoades and Dempsey 1966). According to the authors, once close to these knobs, recombination in adjacent regions was suppressed, influencing such traits. More recently, through fluorescent *in situ* hybridization (FISH), it was demonstrated that knobs are located in areas dense in genes, where large knobs can reduce recombination locally (Ghaffari et al., 2013). Comparisons between European maize genomes and US Corn Belt revealed variation in their repetitive sequences and gene content. The germplasms were separated by the intensity and position of knob regions. However, additional sites with small arrangements of knobs conserved in flint and dent lines were observed. This study also showed that the knob sequences could affect genes surrounded by them, decreasing their expression level (Haberer et al., 2020)

In our study, the heterochromatic knob mapped on the short arm of chromosome 9 was strongly associated with early flowering time. Flowering time-related QTLs have been found across the maize genome (Chardon et al., 2004; Yang et al., 2019). In chromosome 9, flowering time-related QTLs were found, and some genes were identified (Hung et al., 2012; Huang et al., 2017). The main gene located in chromosome 9 was ZmCCT9 (photoperiod sensitive) and mapped on the long arm, opposite to the knob position. Other candidate genes were found along this chromosome, but the precise location was not defined yet (Miller et al., 2008; Li et al., 2016). Further studies should be carried out to map a gene influencing flowering time near the knob on the short arm of chromosome 9.

Constitutive heterochromatins have well-defined roles in the genomes, such as centromeres and telomeres (Salvi et al., 2007; Tacconi and Tarsounas, 2015). That leads us to infer that heterochromatic knobs may also have a role in the maize genome, affecting certain phenotypic traits. However, further investigation in these regions will be needed to find possible candidate genes associated with knobs due to the inhibition of crossing over on their neighborhood.

### CONCLUDING REMARKS

Despite more than 100 years of studies on maize genetics, the discussion on the role of knobs remains a current issue and has shown that this region is an essential fraction of the maize genome. Elimination dynamics of components of the maize genome over successive self-fertilizations were reported (Roessler et al., 2019).Among the five components analyzed in the genome (gene content, ribosomal DNA, B chromosomes, TEs, and knobs), TEs were the most significantly lost components. The results provide insights into the constitutive role played by knobs in the maize genome. Like TEs, knobs are repetitive sequences, which make up about 10% of the maize genome and, for unknown reasons, are not totally eliminated after generations of self-fertilization. That strengthens the hypothesis that heterochromatic knobs may have a functional role within the maize genome, even composed almost exclusively by repetitive sequences. It is interesting to note that over nine cycles of self-fertilizations, the lines used in the present study did not lose all the knobs and even some lines of the JD-4 family conserved all of them.

Throughout the maize genome evolution, it was proposed that flowering time was a trait influenced by changes in the genome size. At high altitudes, maize flowering time was shorter than at low altitudes, followed by a smaller genome (Rayburn et al., 1985, 1994; Poggio et al., 1998; Realini et al., 2015). At the same time, most studies have also indicated a positive correlation between the genome size and the knob abundance (Rayburn et al., 1994; Jian et al., 2017).

We found no significant association between knobs and genome size from our data. The analyses of adapted GWAS carried out showed the contribution of a single locus for the early maize flowering time: the knob present in the short arm of chromosome 9 was associated with reduced flowering time when homozygous or heterozygous. Our results suggest a role of the knobs in the flowering time, different from those previously described. For the first time, maize inbred lines were selected with knobs in specific locations, and their hybrids were developed to carry heterozygous and homozygous knobs. Although the lines have a common origin, they differ in knob composition and in the response to culture *in vitro* (Fluminhan and Aguiar-Perecin, 1998).

The maize flowering time is a complex trait, and several studies have provided insights into its genetic architecture. To this complexity, we add our data that suggest that gene-*like* components, such as specific heterochromatic knobs, are capable of affecting the maize flowering time, probably by interaction with particular genes playing a central role in the regulation of the feature expression. Furthermore, as knobs suppress local recombination (Ghaffari et al., 2013) 6), some genes would be associated with them.

Unravel the mystery behind the repetitive sequences has been a hard work all over the years. It becomes even more difficult for maize since its genome comprises more than 85% of these sequences (Schnable et al., 2009).

We could infer that there would be an interaction between genes controlling flowering time, knob composition and genome size. In addition, transposable elements also contribute to genome size. So, if populations with high content of larger knobs are compared with populations with few smaller knobs, perhaps the effect of knobs on flowering time would be detected, if the effect of genes controlling flowering time were not higher.

## ACKNOWLEDGMENTS

Coordenação de Aperfeiçoamento de Pessoal de Nível Superior— Brasil (CAPES) and Conselho Nacional de Desenvolvimento Científico e Tecnológico (CNPq) for the financial support.

## CONTRIBUTION TO THE FIELD STATEMENT

Maize flowering time is an important agronomic trait for genetic improvement. This feature has been associated with variations in the genome size and heterochromatic knobs. Knobs are constitutive heterochromatin composed of two satellite DNA sequences, 180-pb and TR-1. To show the influence of knob composition in genome size and flowering time, we used sister inbred lines differing in their knob content and originated from a unique plant. Hybrids homozygous and heterozygous for knobs were also obtained from the crossing of some of these lines. The genome size and flowering time of these materials were measured and correlations between knob composition, genome size and flowering time were not observed. Then, an association study was developed and identified a knob marker on chromosome 9 showing a strong association with flowering time. Indeed, modelling allele substitution and dominance effects could offer only one heterochromatic knob locus that could affect flowering time, making it earlier rather than the knob composition.

## AUTHOR CONTRIBUTIONS

MLRAP and MM designed the idea; MLRAP,MM and RFC designed the experiments, RFN designed and carried out statistical analysis; RFC and WRC performed flow cytometry analysis; RFC carried out cytogenetic analysis, field experiments and managed the results; RFC, MM and MLRAP wrote the paper.

## Supplementary Materials

### Figure Captions

**Figure S1. Flowering time and Genome Size versus knob condition**. Boxplots show the comparison between male and female flowering time (days) and genome size (picograms) regarding heterozygous knobs or homozygous knobs. The boxes indicate the first quartile (lower line), the second quartile or mean (central line), and the third quartile (upper line). Additionally, the whiskers represent the standard deviation with the dots as the outliers.

**Figure S2. QQ-plot and Manhattan plot of GWAS for Genome size**. Plots characterize the allele substitution effect. Significant associations are not found. The x axis shows the knobs K3L, K5L, K7S and K9S. The y axis shows the logarithm of the value of significance.

**Figure S3. QQ-plot of GWAS of flowering time**. Plots characterize the allele substitution effect for **(A)** male flowering (mf) and **(B)** female flowering (ff) time. The dots represent the knobs K3L, K5L, K7S and K9S, respectively.

**Figure S4. QQ-plot of GWAS of flowering time**. Plots characterize the dominance effect for **(A)** male flowering (mf) and **(B)** female flowering (ff) time. The dots represent the knobs K3L, K5L, K7S and K9S respectively.

