## Supplementary figures and images for "A heterochromatic knob reducing the flowering time in maize"

### Figure S1

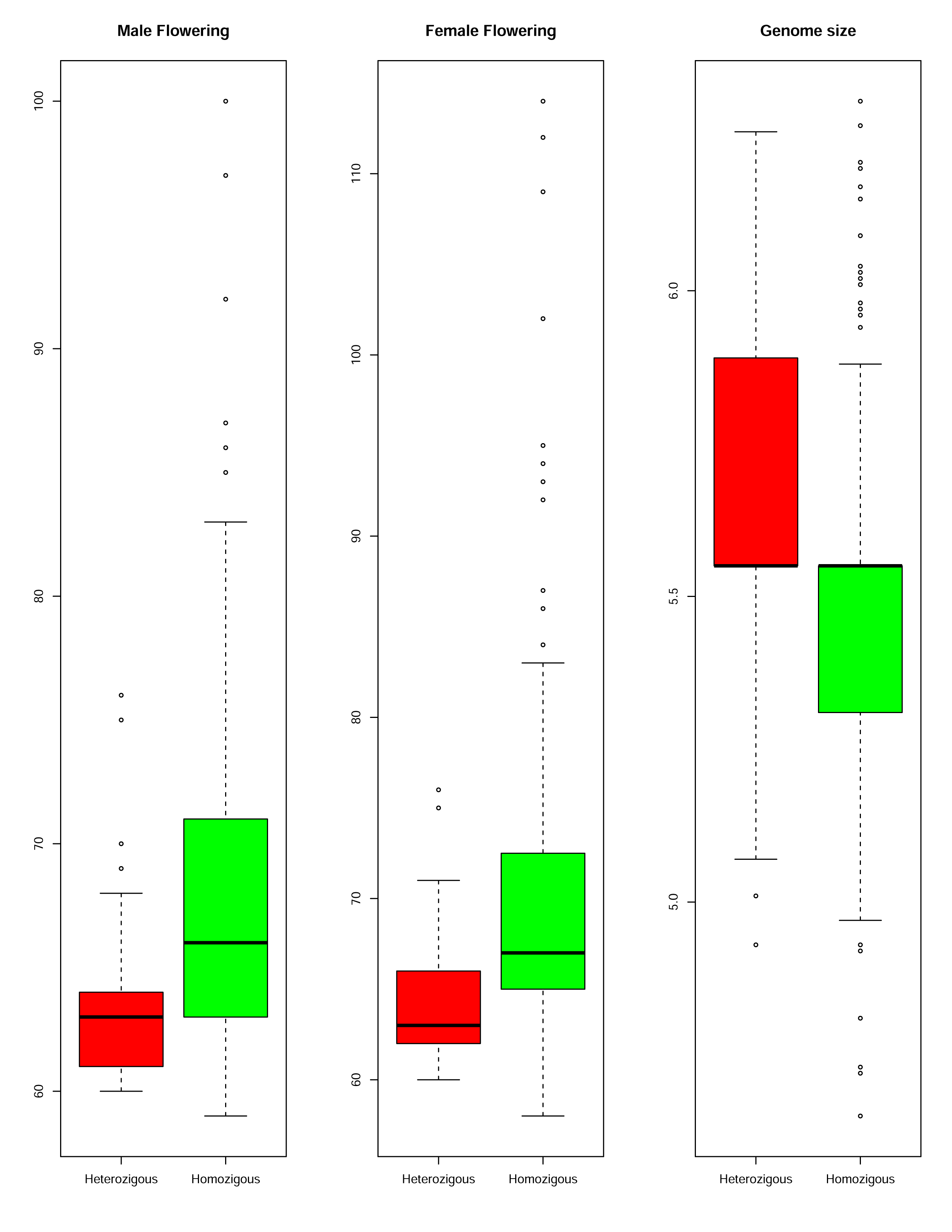

### Figure S2

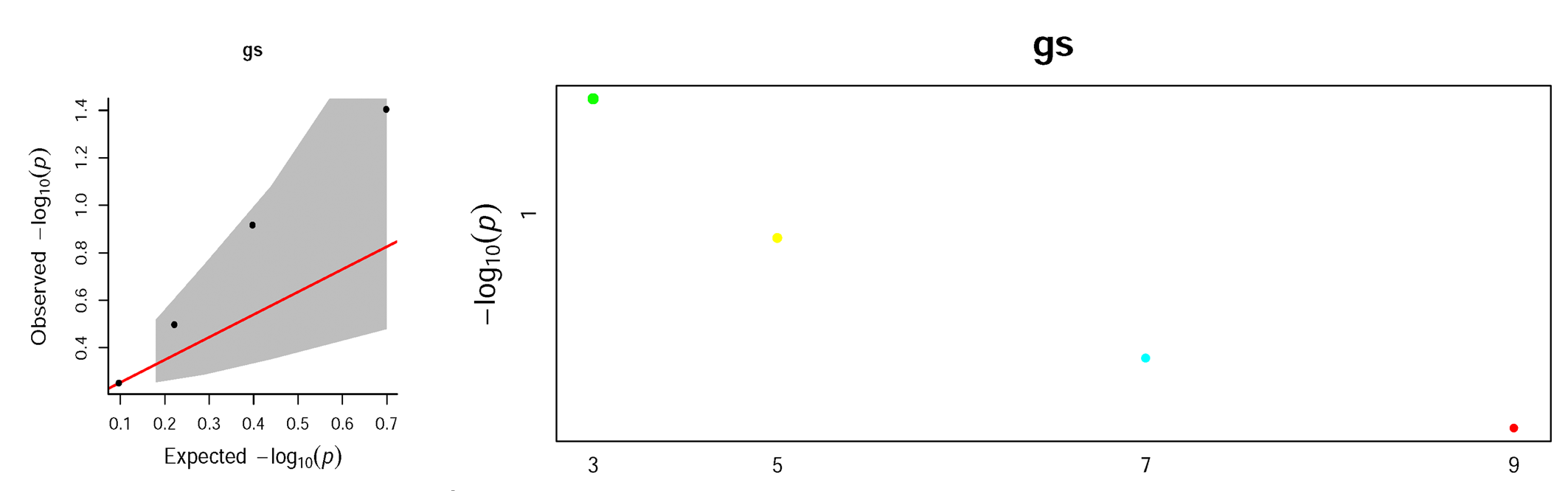

### Figure S3

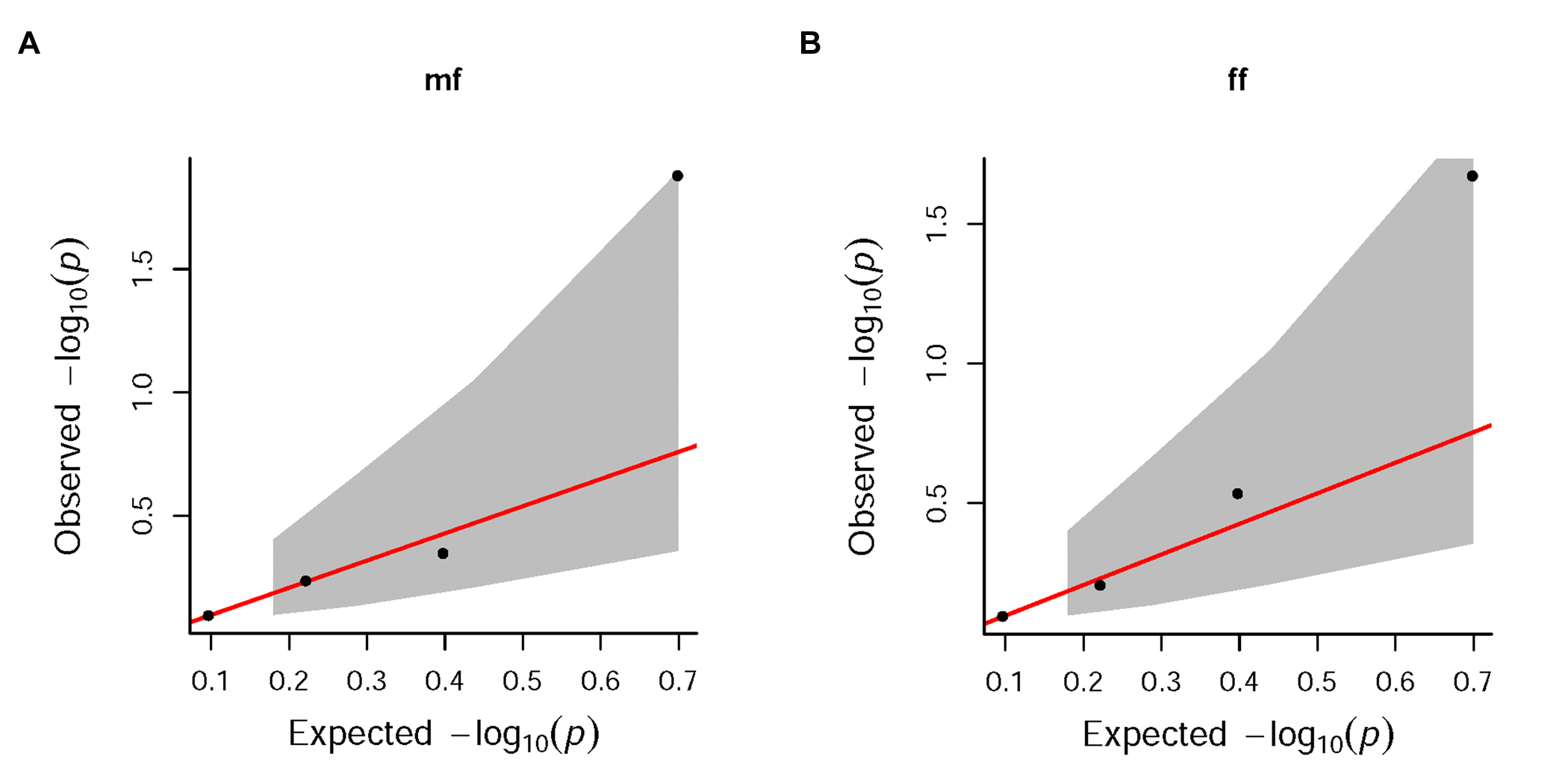

### Figure S4

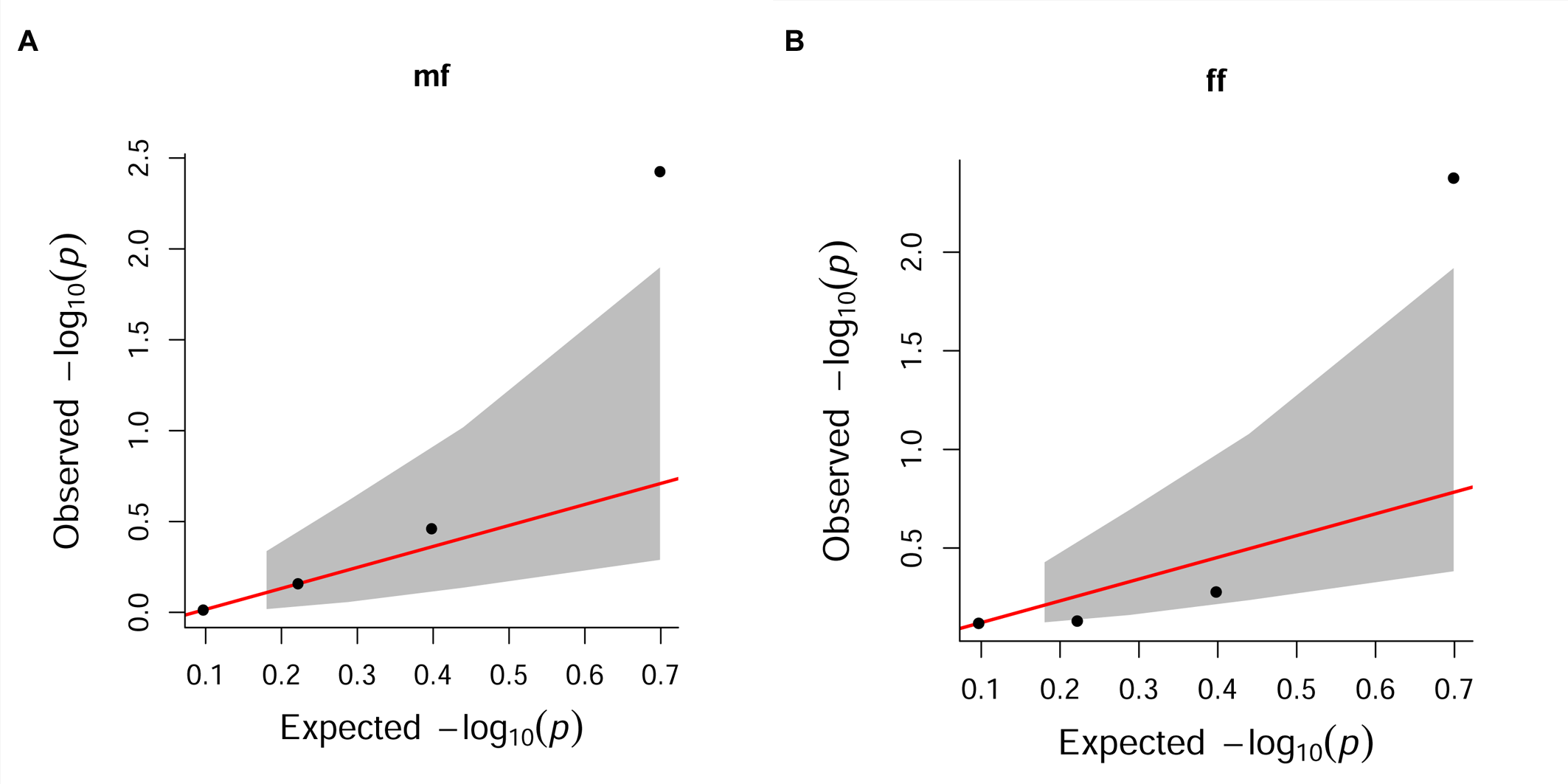
