## Supplementary material for "A heterochromatic knob reducing the flowering time in maize": Table S1

**Table S1**. The origin of the inbred lines from the Jac Duro variety: upper the table shows the original knob composition, and the bottom shows the selected inbred lines and their current knob composition in S9.

| **Jac Duro Original - Cateto Variety** | | | | | | | |
| --- | --- | --- | --- | --- | --- | --- | --- |
| **K3L** | **K5L** | **K6L1/K6L1** | **K7S** | **K7L** | **K8L1/K8L2** | | **K9S** |
| segregating | segregating | homozygous knobs | segregating | homozygous knobs | homozygous knobs | | segregating |
| ++/+0/00 | ++/+0/00 | ++ | ++/+0/00 | ++ | ++ | | ++/+0/00 |
|  | **Knob** | | | | |  | |
| **Genotype** | **K3L** | **K5L** | **K7S** | **K9S** | **Generation** | |  |
| 99 | - | - | - | - | S1 | |  |
| 200-14 | - | - | - | - | S2 | |  |
| 240-14-4 | ++/+0/00 | ++/+0/00 | ++/+0/00 | ++/+0/00 | S2 | |  |
| 240-14-1 | ++/+0/00 | 00 | ++ | ++/+0/00 | S3 | |  |
| 240-14-2 | ++ | 00 | 00 | ++/+0/00 | S3 | |  |
| 4-1 | ++/+0/00 | ++/+0/00 | ++/+0/00 | ++/+0/00 | S3 | |  |
| 4-4 | ++/+0/00 | ++/+0/00 | ++/+0/00 | ++/+0/00 | S3 | |  |
| 1-1 | - | - | - | - | S4 | |  |
| 1-2 | - | - | - | - | S4 | |  |
| 1-3 | - | - | - | - | S4 | |  |
| 2-1 | - | - | - | - | S4 | |  |
| 2-2 | - | - | - | - | S4 | |  |
| 2-3 | - | - | - | - | S4 | |  |
| 41-1 | +0/00 | ++ | 00 | ++/+0/00 | S4 | |  |
| 41-2 | ++/+0/00 | ++ | 00 | ++ | S4 | |  |
| 44-1 | ++/+0/00 | ++/00 | ++/+0/00 | ++/+0/00 | S4 | |  |
| 44-2 | ++/+0 | ++/+0/00 | ++/00 | ++/+0/00 | S4 | |  |
| 44-4 | 00 | ++ | ++/+0/00 | +0/00 | S4 | |  |
| 13-1 | 00 | 00 | ++ | ++ | S5 | |  |
| 13-2 | 00 | 00 | ++ | ++ | S5 | |  |
| 13-3 | ++/+0/00 | 00 | ++ | ++ | S5 | |  |
| 21-1 | ++ | 00 | 00 | ++ | S5 | |  |
| 21-3 | ++ | 00 | 00 | ++ | S5 | |  |
| 411-2 | ++ | ++ | 00 | ++/00 | S5 | |  |
| 412-3 | ++ | ++/+0 | 00 | ++/+0 | S5 | |  |
| 412-4 | ++/+0/00 | ++/+0 | 00 | ++/+0 | S5 | |  |
| 441-1 | ++ | ++/+0 | ++ | ++ | S5 | |  |
| 441-3 | ++ | ++ | ++/+0/00 | ++/+0 | S5 | |  |
| 442-2 | +0 | ++/+0 | ++/+0/00 | ++ | S5 | |  |
| 444-3 | 00 | ++ | 00 | 00 | S5 | |  |
| 131-1 | 00 | 00 | ++ | ++ | S6 | |  |
| 131-5 | 00 | 00 | ++ | ++ | S6 | |  |
| 132-3 | 00 | 00 | ++ | ++ | S6 | |  |
| 133-4 | +0/00 | 00 | ++ | 00 | S6 | |  |
| 211-1 | ++ | 00 | 00 | ++ | S6 | |  |
| 211-2 | ++ | 00 | 00 | ++ | S6 | |  |
| 213-1 | ++ | 00 | 00 | ++ | S6 | |  |
| 213-3 | ++ | 00 | 00 | ++ | S6 | |  |
| 4112-1 | ++ | ++ | 00 | ++ | S6 | |  |
| 4112-2 | ++ | ++ | 00 | ++ | S6 | |  |
| 4123-3 | ++ | ++ | 00 | ++ | S6 | |  |
| 4123-4 | ++ | ++ | 00 | ++ | S6 | |  |
| 4124-2 | 00 | ++ | 00 | ++ | S6 | |  |
| 4411-2 | ++ | ++ | ++ | ++ | S6 | |  |
| 4411-3 | ++ | ++ | ++ | ++ | S6 | |  |
| 4413-1 | ++ | ++ | ++/00 | ++ | S6 | |  |
| 4413-2 | ++ | ++ | ++/+0/00 | ++ | S6 | |  |
| 4422-4 | ++/+0/00 | ++/+0 | 00 | ++ | S6 | |  |
| 4426-1 | ++ | ++ | ++ | 00 | S6 | |  |
| 4443-1 | 00 | ++ | 00 | 00 | S6 | |  |
| 1311-1 | 00 | 00 | ++ | ++ | S7 | |  |
| 1315-1 | 00 | 00 | ++ | ++ | S7 | |  |
| 1323-3 | 00 | 00 | ++ | ++ | S7 | |  |
| 1334-2 | +0/00 | 00 | ++ | 00 | S7 | |  |
| 2111-2 | ++ | 00 | 00 | ++ | S7 | |  |
| 2111-3 | ++ | 00 | 00 | ++ | S7 | |  |
| 2112-1 | ++ | 00 | 00 | ++ | S7 | |  |
| 21311-2 | ++ | 00 | 00 | ++ | S7 | |  |
| 21331-1 | ++ | 00 | 00 | ++ | S7 | |  |
| 44433-2 | 00 | ++ | 00 | 00 | S7 | |  |
| 13131-1 | 00 | 00 | ++ | ++ | S8 | |  |
| 13151-1 | 00 | 00 | ++ | ++ | S8 | |  |
| 13233-1 | 00 | 00 | ++ | ++ | S8 | |  |
| 13342-1 | ++ | 00 | ++ | 00 | S8 | |  |
| 13342-2 | ++ | 00 | ++ | 00 | S8 | |  |
| 13342-7 | 00 | 00 | ++ | 00 | S8 | |  |
| 21112-1 | ++ | 00 | 00 | ++ | S8 | |  |
| 21113-1 | ++ | 00 | 00 | ++ | S8 | |  |
| 21121-1 | ++ | 00 | 00 | ++ | S8 | |  |
| 21311-2 | ++ | 00 | 00 | ++ | S8 | |  |
| 21331-1 | ++ | 00 | 00 | ++ | S8 | |  |
| 441123-3 | ++ | ++ | ++ | ++ | S8 | |  |
| 441132-2 | ++ | ++ | ++ | ++ | S8 | |  |
| 441311-2 | ++ | ++ | ++/+0/00 | ++ | S8 | |  |
| 441324-1 | ++ | ++ | ++ | ++ | S8 | |  |
| 442213-1 | ++ | ++ | ++ | ++ | S8 | |  |
| 442213-2 | 00 | ++ | ++ | ++ | S8 | |  |
| 442242-1 | 00 | ++ | +0/00 | ++ | S8 | |  |
| 442612-1 | 00 | ++ | ++ | 00 | S8 | |  |
| 444332 | 00 | ++ | 00 | 00 | S8 | |  |
| 131311-1 | 00 | 00 | ++ | ++ | S9 | |  |
| 131511 | 00 | 00 | ++ | ++ | S9 | |  |
| 132331-1 | 00 | 00 | ++ | ++ | S9 | |  |
| 133421 | ++ | 00 | ++ | 00 | S9 | |  |
| 133422 | ++ | 00 | ++ | 00 | S9 | |  |
| 133425 | 00 | 00 | ++ | 00 | S9 | |  |
| 133427 | 00 | 00 | ++ | 00 | S9 | |  |
| 211121 | ++ | 00 | 00 | ++ | S9 | |  |
| 211131 | ++ | 00 | 00 | ++ | S9 | |  |
| 211211 | ++ | 00 | 00 | ++ | S9 | |  |
| 213112 | ++ | 00 | 00 | ++ | S9 | |  |
| 213311 | ++ | 00 | 00 | ++ | S9 | |  |
| 441123-3 | ++ | ++ | ++ | ++ | S9 | |  |
| 441132-2 | ++ | ++ | ++ | ++ | S9 | |  |
| 441311-2 | ++ | ++ | ++ | ++ | S9 | |  |
| 441324-1 | ++ | ++ | ++ | ++ | S9 | |  |
| 442213-1 | ++ | ++ | ++ | ++ | S9 | |  |
| 442213-2 | 00 | ++ | ++ | ++ | S9 | |  |
| 442242-1 | 00 | ++ | 00 | ++ | S9 | |  |
| 442612-1 | 00 | ++ | ++ | 00 | S9 | |  |
