## Supplementary material for "A heterochromatic knob reducing the flowering time in maize": Table S2

**Table S2.** Average values of phenotypic measurements and the cytological knob mapping; Genotype, Genotype ID (Gid), Maternal parent (Female), Paternal parent (Male), Material type (hybrid/line), Year (2018/2019), Knob condition (homozygous/heterozygous), Knob position* (K3L/K5L/K7S/K9S), Genome size (pg and Mbp), Female flowering (ff), Male flowering (mf).

|  | | | | | | | | Knob |  |  | Genome size |  | |
| --- | --- | --- | --- | --- | --- | --- | --- | --- | --- | --- | --- | --- | --- |
| Genotype | Gid | Female | Male | Type | Year | Condition | 3L | 5L | 7S | 9S | pg Mbp** | ff | mf |
| 44431-04 | Gen 36 | 44431-04 | 44431-04 | line | 2018 | homozigous | -1 | 1 | -1 | -1 | NA NA | 68 | 69.33 |
| 442242/1-04 | Gen 37 | 442242/1-04 | 442242/1-04 | line | 2018 | homozigous | -1 | 1 | -1 | 1 | NA NA | 65.33 | 64.33 |
| 442612/1-04 | Gen 38 | 442612/1-04 | 442612/1-04 | line | 2018 | homozigous | -1 | 1 | -1 | 1 | NA NA | 64 | 62 |
| 441311/2-04 | Gen 39 | 441311/2-04 | 441311/2-04 | line | 2018 | homozigous | 1 | 1 | 1 | 1 | NA NA | 66.33 | 64.67 |
| 441324/1-04 | Gen 40 | 441324/1-04 | 441324/1-04 | line | 2018 | homozigous | 1 | 1 | 1 | 1 | NA NA | 62 | 63 |
| 442213/1 | Gen 41 | 442213/1 | 442213/1 | line | 2018 | homozigous | 1 | 1 | 1 | 1 | 4.97 4860.66 | 68 | 68.33 |
| 44431-04 | Gen 36 | 44431-04 | 44431-04 | line | 2019 | homozigous | -1 | 1 | -1 | -1 | 5.45 5330.1 | 84 | 77.25 |
| 442242/1-04 | Gen 37 | 442242/1-04 | 442242/1-04 | line | 2019 | homozigous | -1 | 1 | -1 | 1 | 5.55 5427.9 | 76 | 75.2 |
| 442612/1-04 | Gen 38 | 442612/1-04 | 442612/1-04 | line | 2019 | homozigous | -1 | 1 | -1 | 1 | 5.49 5369.22 | 66.4 | 66.2 |
| 441311/2-04 | Gen 39 | 441311/2-04 | 441311/2-04 | line | 2019 | homozigous | 1 | 1 | 1 | 1 | 5.3 5183.4 | 70.8 | 67 |
| 441324/1-04 | Gen 40 | 441324/1-04 | 441324/1-04 | line | 2019 | homozigous | 1 | 1 | 1 | 1 | 5.17 5056.26 | 66.2 | 65.4 |
| 441123/1-04 | Gen 53 | 441123/1-04 | 441123/1-04 | line | 2019 | homozigous | 1 | 1 | 1 | 1 | NA NA | 67.8 | 65.2 |
| 441123/3-04 | Gen 54 | 441123/3-04 | 441123/3-04 | line | 2019 | homozigous | 1 | 1 | 1 | 1 | 5.4 5281.2 | 75.2 | 71.6 |
| 441132/2-04 | Gen 55 | 441132/2-04 | 441132/2-04 | line | 2019 | homozigous | 1 | 1 | 1 | 1 | 5.33 5212.74 | 70.2 | 66.6 |
| 444332-04 | Gen 56 | 444332-04 | 444332-04 | line | 2019 | homozigous | -1 | 1 | -1 | -1 | 5.55 5427.9 | 80.6 | 76.4 |
| 131311/1-04 | Gen 42 | 131311/1-04 | 131311/1-04 | line | 2018 | homozigous | -1 | -1 | 1 | 1 | NA NA | 65 | 64 |
| 132331/1-04 | Gen 43 | 132331/1-04 | 132331/1-04 | line | 2018 | homozigous | -1 | -1 | 1 | 1 | NA NA | 65.67 | 64 |
| 131311/1-04 | Gen 42 | 131311/1-04 | 131311/1-04 | line | 2019 | homozigous | -1 | -1 | 1 | 1 | 5.6 5476.8 | 79.2 | 76.4 |
| 132331/1-04 | Gen 43 | 132331/1-04 | 132331/1-04 | line | 2019 | homozigous | -1 | -1 | 1 | 1 | 5.49 5369.22 | 75.2 | 71.4 |

Knob	Genome size

| Genotype | Gid | Female | Male | Type | Year | Condition | 3L | 5L | 7S | 9S | pg | Mbp** | ff | mf |
| --- | --- | --- | --- | --- | --- | --- | --- | --- | --- | --- | --- | --- | --- | --- |
| 131511-04 | Gen 44 | 131511-04 | 131511-04 | line | 2019 | homozigous | -1 | -1 | 1 | 1 | NA | NA | 71.8 | 67.8 |
| 133421-04 | Gen 45 | 133421-04 | 133421-04 | line | 2019 | homozigous | 1 | -1 | 1 | -1 | 5.49 | 5369.22 | 66.2 | 65.4 |
| 133422-04 | Gen 46 | 133422-04 | 133422-04 | line | 2019 | homozigous | 1 | -1 | 1 | -1 | 5.51 | 5388.78 | 90 | 85.8 |
| 133427-04 | Gen 47 | 133427-04 | 133427-04 | line | 2019 | homozigous | -1 | -1 | 1 | -1 | 5.55 | 5427.9 | 76.6 | 70.4 |
| 211121/1-04 | Gen 48 | 211121/1-04 | 211121/1-04 | line | 2019 | homozigous | 1 | -1 | -1 | 1 | 5.55 | 5427.9 | 84.2 | 75.8 |
| 211131-04 | Gen 49 | 211131-04 | 211131-04 | line | 2019 | homozigous | 1 | -1 | -1 | 1 | NA | NA | 70 | 70 |
| 211211/2-04 | Gen 50 | 211211/2-04 | 211211/2-04 | line | 2019 | homozigous | 1 | -1 | -1 | 1 | 5.42 | 5300.76 | 74.6 | 73.6 |
| 213112-04-04 | Gen 51 | 213112-04-04 | 213112-04-04 | line | 2019 | homozigous | 1 | -1 | -1 | 1 | 5.47 | 5349.66 | 66.6 | 67.4 |
| 213311/1 | Gen 52 | 213311/1 | 213311/1 | line | 2019 | homozigous | 1 | -1 | -1 | 1 | 5.18 | 5066.04 | 66.6 | 65.4 |
| 44431x442242/1 | Gen 1 | 44431 | 442242/1 | hybrid | 2018 | heterozigous | -1 | 1 | -1 | 0 | 5.62 | 5496.36 | 64 | 62.6 |
| 44431x442612/1 | Gen 2 | 44431 | 442612/1 | hybrid | 2018 | heterozigous | -1 | 1 | -1 | 0 | 5.78 | 5652.84 | 61.6 | 61.2 |
| 44431x441311/1 | Gen 3 | 44431 | 441311/1 | hybrid | 2018 | heterozigous | 0 | 1 | 0 | 0 | 5.36 | 5242.08 | 64.6 | 63.8 |
| 44431x441324/1 | Gen 4 | 44431 | 441324/1 | hybrid | 2018 | heterozigous | 0 | 1 | 0 | 0 | 5.55 | 5427.9 | 63.5 | 63 |
| 44431x442213/1 | Gen 5 | 44431 | 442213/1 | hybrid | 2018 | heterozigous | 0 | 1 | 0 | 0 | 5.43 | 5310.54 | 66 | 64.2 |
| 442612/1x44431 | Gen 10 | 442612/1 | 44431 | hybrid | 2018 | heterozigous | -1 | 1 | -1 | 0 | 5.64 | 5515.92 | 64.4 | 62.6 |
| 442612/1x442242/1 | Gen 11 | 442612/1 | 442242/1 | hybrid | 2018 | homozigous | -1 | 1 | -1 | 1 | NA | NA | 69 | 65.33 |
| 442612/1x441311/2 | Gen 12 | 442612/1 | 441311/2 | hybrid | 2018 | heterozigous | 0 | 1 | 0 | 1 | 5.55 | 5427.9 | 65.4 | 64 |
| 442612/1x441324/1 | Gen 13 | 442612/1 | 441324/1 | hybrid | 2018 | heterozigous | 0 | 1 | 0 | 1 | 5.84 | 5711.52 | 68 | 65.66 |
| 442612/1x442213/1 | Gen 14 | 442612/1 | 442213/1 | hybrid | 2018 | heterozigous | 0 | 1 | 0 | 1 | NA | NA | 66.5 | 65 |
| 442612/1x131311/1 | Gen 15 | 442612/1 | 131311/1 | hybrid | 2018 | heterozigous | 0 | 1 | 0 | 1 | 5.77 | 5643.06 | 65.25 | 63.5 |
| 441311/2x44431 | Gen 16 | 441311/2 | 44431 | hybrid | 2018 | heterozigous | 0 | 1 | 0 | 0 | 5.48 | 5359.44 | 62.8 | 61.6 |
| 441311/1x442242/1 | Gen 17 | 441311/1 | 442242/1 | hybrid | 2018 | heterozigous | 0 | 1 | 0 | 1 | 5.64 | 5515.92 | 63.8 | 62 |
| 441311/2x442612/1 | Gen 18 | 441311/2 | 442612/1 | hybrid | 2018 | heterozigous | 0 | 1 | 0 | 1 | 5.89 | 5760.42 | 63.6 | 62.6 |
| 441311/2x441324/1 | Gen 19 | 441311/2 | 441324/1 | hybrid | 2018 | homozigous | 1 | 1 | 1 | 1 | 5.35 | 5232.3 | 67 | 65.4 |
| 441311/2x442213/1 | Gen 20 | 441311/2 | 442213/1 | hybrid | 2018 | homozigous | 1 | 1 | 1 | 1 | 5.82 | 5691.96 | 65.2 | 62.4 |
| 441324/1x44431 | Gen 21 | 441324/1 | 44431 | hybrid | 2018 | heterozigous | 0 | 1 | 0 | 0 | 5.54 | 5418.12 | 62.2 | 62.4 |

Knob	Genome size

| Genotype | Gid | Female | Male | Type | Year | Condition | 3L | 5L | 7S | 9S | pg | Mbp** | ff | mf |
| --- | --- | --- | --- | --- | --- | --- | --- | --- | --- | --- | --- | --- | --- | --- |
| 441324/1x442241/1 | Gen 22 | 441324/1 | 442241/1 | hybrid | 2018 | heterozigous | 0 | 1 | 0 | 1 | 5.59 | 5467.02 | 64 | 62.4 |
| 441324/1x442612/1 | Gen 23 | 441324/1 | 442612/1 | hybrid | 2018 | heterozigous | 0 | 1 | 0 | 1 | 5.47 | 5349.66 | 63.8 | 63.2 |
| 441324/1x441311/2 | Gen 24 | 441324/1 | 441311/2 | hybrid | 2018 | homozigous | 1 | 1 | 1 | 1 | NA | NA | 64 | 60.75 |
| 441324/1x442213/1 | Gen 25 | 441324/1 | 442213/1 | hybrid | 2018 | homozigous | 1 | 1 | 1 | 1 | 5.64 | 5515.92 | 65.6 | 65 |
| 441324/1x131311/1 | Gen 26 | 441324/1 | 131311/1 | hybrid | 2018 | heterozigous | 0 | 0 | 1 | 1 | 5.94 | 5809.32 | 62.6 | 60.8 |
| 442213/1x4443/1 | Gen 27 | 442213/1 | 4443/1 | hybrid | 2018 | heterozigous | 0 | 1 | 0 | 0 | 5.34 | 5222.52 | 64.6 | 64 |
| 442213/1x442242/1 | Gen 28 | 442213/1 | 442242/1 | hybrid | 2018 | heterozigous | 0 | 1 | 0 | 1 | NA | NA | 68.6 | 67.8 |
| 442213/1x442612/1 | Gen 29 | 442213/1 | 442612/1 | hybrid | 2018 | heterozigous | 0 | 1 | 0 | 1 | 5.41 | 5290.98 | 68.25 | 67.75 |
| 442213/1x441311/2 | Gen 30 | 442213/1 | 441311/2 | hybrid | 2018 | homozigous | 1 | 1 | 1 | 1 | 5.8 | 5672.4 | 66.8 | 63 |
| 442213/1x441324/1 | Gen 31 | 442213/1 | 441324/1 | hybrid | 2018 | homozigous | 1 | 1 | 1 | 1 | 5.65 | 5525.7 | 62.6 | 62 |
| 442242/1x44431 | Gen 6 | 442242/1 | 44431 | hybrid | 2018 | heterozigous | -1 | 1 | -1 | 0 | 5.55 | 5427.9 | 63.8 | 61.8 |
| 442242/1x442612/1 | Gen 7 | 442242/1 | 442612/1 | hybrid | 2018 | homozigous | -1 | 1 | -1 | 1 | 5.89 | 5760.42 | 67 | 66 |
| 442242/1 x 441311/2 | Gen 8 | 442242/1 | 441311/2 | hybrid | 2018 | heterozigous | 0 | 1 | 0 | 1 | 4.93 | 4821.54 | 63 | 62 |
| 442242/1x441324/1 | Gen 9 | 442242/1 | 441324/1 | hybrid | 2018 | heterozigous | 0 | 1 | 0 | 1 | 5.58 | 5457.24 | 62 | 60 |
| 131311/1x442612/1 | Gen 32 | 131311/1 | 442612/1 | hybrid | 2018 | heterozigous | -1 | 0 | 0 | 1 | 5.64 | 5515.92 | 62 | 61 |
| 131311/1x441324/1 | Gen 33 | 131311/1 | 441324/1 | hybrid | 2018 | heterozigous | 0 | 0 | 1 | 1 | 5.37 | 5251.86 | 63.2 | 61.6 |
| 131311/1x132331/1 | Gen 34 | 131311/1 | 132331/1 | hybrid | 2018 | homozigous | -1 | -1 | 1 | 1 | 5.85 | 5721.3 | 62.6 | 62.4 |
| 132331/1x131311/1 | Gen 35 | 132331/1 | 131311/1 | hybrid | 2018 | homozigous | -1 | -1 | 1 | 1 | 5.05 | 4938.9 | 64.2 | 63.4 |

* Knob position matrix: -1 = knob absence, 0 = heterozygous knob presence and 1 = homozygous knob presence. NA, non analyzed.

** 1pg = 0.978 x 10^9^ bp (Reference: J. Doležel, J. Greilhuber, J. Suda, Nuclear DNA content and genome size of trout and human. Cytometry Part A. **51A**, 127–128 (2003)).
