## Supplementary material for "A heterochromatic knob reducing the flowering time in maize": Table S3

**Table S3. List of additive and dominance effects for flowering time, genome size, and knob positions.** Male flowering (MF), female flowering (FF), genome size (GS), Knob position, p-value via Bonferroni test, and minor allele frequency (MAF)

| **Trait Marker** | **Chr.** | **Position** | **MAF** | **Additive**  **Effect** | ***P-value*** | **Dominance**  **Effect** | ***P-value*** |
| --- | --- | --- | --- | --- | --- | --- | --- |
| K3L | 3 | Long arm | 0.48 | 0.13 | 0.58 | 0.40 | 0.35 |
| MF K5L | 5 | Long arm | 0.26 | -0.11 | 0.45 | 0.01 | 0.97 |
| K7S | 7 | Short arm | 0.44 | 0.06 | 0.81 | 0.17 | 0.70 |
| K3L | 3 | Long arm | 0.48 | -0.05 | 0.81 | 0.74 | 0.13 |
| FF K5L | 5 | Long arm | 0.26 | -0.07 | 0.62 | 0.76 | 0.10 |
| K7S | 7 | Short arm | 0.44 | 0.25 | 0.29 | 0.52 | 0.25 |
| K3L | 3 | Long arm | 0.48 | 0.01 | 0.04 | -0.01 | 0.03 |
| GS K5L | 5 | Long arm | 0.26 | 0.004 | 0.12 | -0.01 | 0.23 |
| K7S | 7 | Short arm | 0.44 | 0.005 | 0.32 | -0.01 | 0.52 |
